# Structure and mechanism of the RalGAP tumor suppressor complex

**DOI:** 10.1101/2024.11.25.625123

**Authors:** René Rasche, Björn Udo Klink, Lisa Helene Apken, Esther Michalke, Minghao Chen, Andrea Oeckinghaus, Christos Gatsogiannis, Daniel Kümmel

## Abstract

The RalGAP (GTPase activating protein) complexes are negative regulators of the Ral GTPases and thus crucial components that counteract (oncogenic) Ras signaling. However, no structural information on the architecture of this tumor suppressor complex is available hampering a mechanistic understanding of its functionality. Here, we present a cryo-EM structure of RalGAP that reveals an extended 58 nm tetrameric architecture comprising two heterodimers of the RalGAPα and RalGAPβ subunits. We show that the catalytic domain of RalGAPα requires stabilization by a unique domain of RalGAPβ, providing the molecular basis for why RalGAP complexes are obligatory heterodimers. Formation of RalGAP tetramers is not required for activity *in vitro*, but essential for function of the complex *in vivo*. Structural analysis of RalGAP subunit variants reported in cancer patients suggests effects on complex formation and thus functional relevance in tumor development, emphasizing the significance of the obtained structural information for medical research.

## Introduction

The Ral GTPases RalA and RalB act as molecular switches that regulate vesicular trafficking and signal transduction, thereby impacting exocytosis, endocytosis, cell survival and proliferation^1^. The best studied Ral effectors are the exocyst subunits Sec5 and Exo84^2–4^ and the Ral binding protein 1 (RalBP1, or RLIP76)^5–7^, which all three provide a link to membrane transport. Via Sec5, RalB has also been shown to activate Tank-binding kinase 1 (TBK1) in an exocyst-independent manner to promote tumor cell survival^8^. Importantly, the Ral signaling pathway has long been recognized for its crucial function downstream of the Ras GTPases^9^. Rals have been shown to support Ras-dependent tumorigenesis but have also been implicated in cancer progression independent of Ras^1^.

The RalA and RalB isoforms are 82% identical and belong to the Ras superfamily of small GTPases. The activation of Rals from a GDP-bound ‘off’ state to a GTP-bound ‘on’ state is mediated by guanine nucleotide exchange factors (GEFs), some of which are directly controlled by the Ras GTPases. The two RalGAP complexes, RalGAP1 and RalGAP2, act as GAPs (GTPase activating proteins) that limit Ral-GTP levels by promoting hydrolytic activity. They comprise one of the catalytic α-subunits RalGAPα1 (RGα1) in the RalGAP1 complex or RalGAPα2 (RGα2) in theRalGAP2 complex, and a common regulatory β-subunit (RGβ)^10–12^ (Fig 1A). Both RGα and RGβ contain asparagine GAP(-like) domains. First identified in RapGAP, this type of GAP domain uses a unique catalytic mechanism by employing a conserved asparagine to position a water in the GTPase nucleotide binding pocket to promote GTP hydrolysis^13–15^. The GAP domain of RGβ lacks the catalytic asparagine residue and is thus not active^10^, but RalGAP complexes require the conserved asparagine of the GAP domain of the RGα subunits for activity. However, RGα alone is not functional and depends on binding of RGβ with the underlying molecular details remaining unclear^10^.

**Figure 1:**
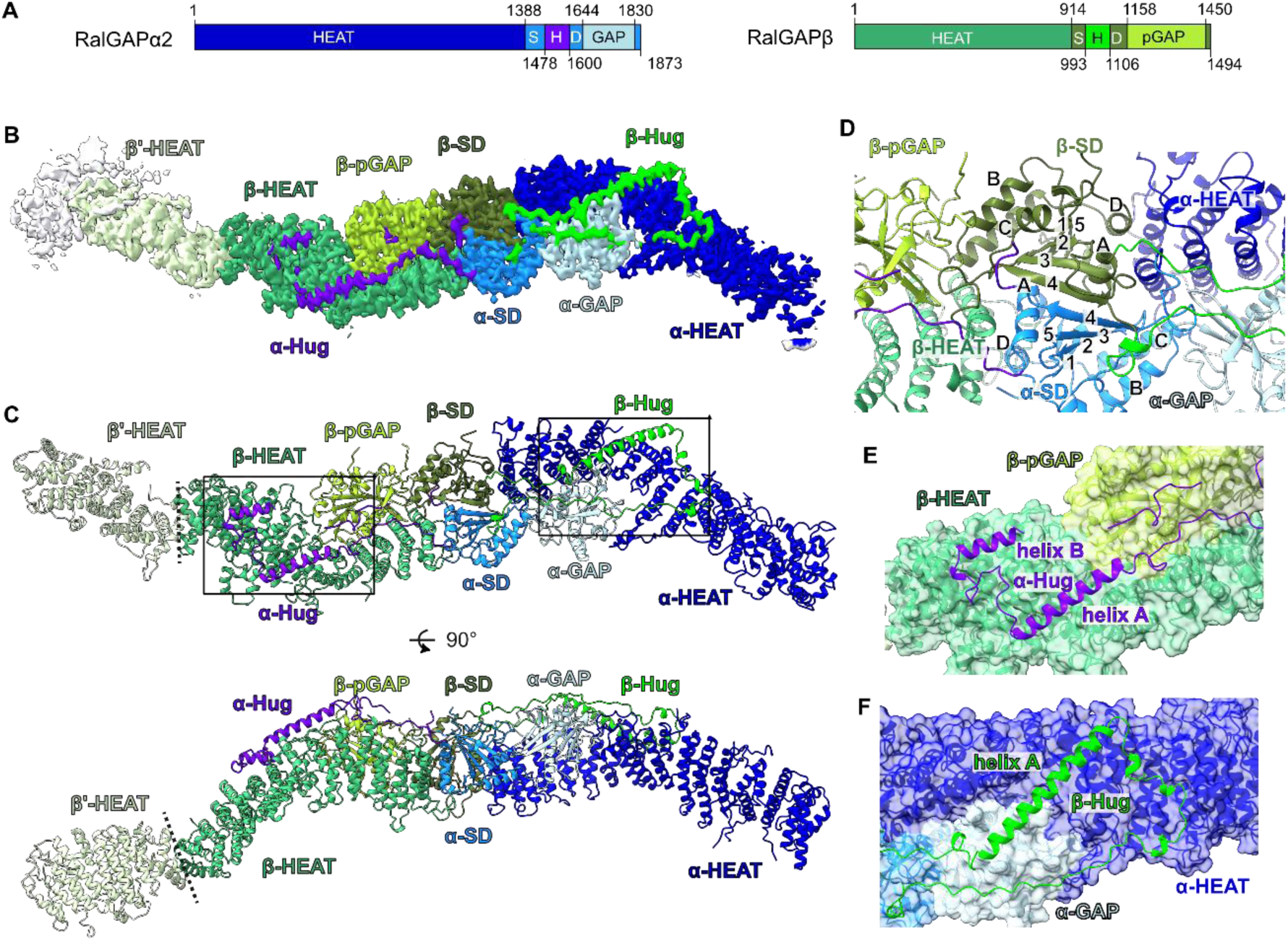
Cryo-EM structure of the RalGAP complex. (A) Domain architecture of RGα2 and RGβ. HEAT: α-solenoid HEAT repeat-like domain; SD: split stabilization and dimerization domain; H: hug domain; (p)GAP: (pseudo-) GTPase activating protein domain. (B) Composite experimental map of RalGAP (half)-particles. (C) Model of the RGα2/RGβ heterodimer and the N-terminal portion of a second RGβ subunit. Domains are colored as in 1A. Dashed lines indicate the interface of both half-particles. (D) Close-up of the SD domain heterodimerization interface. The β-strands (1-5) and α-helices (A-D) of the SD domains are labelled. (E) Interaction of the RGα2 hug domain with RGβ. (F) Interaction of the RGβ hug domain with RGα2.

Another RapGAP domain-containing protein is TSC2 (tuberous sclerosis complex 2), which together with TSC1 and TBC1D7 forms the TSC complex^16–18^. The TSC2 GAP domain accelerates GTP hydrolysis of the GTPase Rheb^15,19–22^. Because Rheb is required for the activation of the mTORC1 (mechanistic target of rapamycin complex 1) kinase^23^, the master regulator of cellular growth, the TSC complex is an important tumor suppressor and loss of function causes the genetic disease TSC^24^.

In addition, the non-canonical κB-Ras GTPases (of which two isoforms, κB-Ras 1 and κB-Ras 2 are present in humans) bind to the N-termini of both RGα subunits and support complex function *in vivo* with the underlying molecular reason remaining elusive^25,26^. Activity of RalGAP can furthermore be negatively regulated by AKT phosphorylation and 14-3-3 protein binding^11,12,27^, which is also reminiscent of findings reported for the TSC complex^28^.

Despite the ability of Ral GTPases to promote tumorigenesis, no Ral mutations with oncogenic potential analogous to the Ras GTPases have been described in patients, and neither has relevance of mutations in Ral effectors or regulators been studied^29^. However, low expression of RalGAPα2 leads to an increase in tumor cell proliferation, invasion and migration in cancer cell lines or mouse models of hepatocellular carcinoma^30^, colitis-associated cancer^31^, oral squamous cell carcinoma^32^, prostate cancer^33^, and bladder cancer^34^. A comprehensive proteomic analysis of 140 patient pancreatic tumor samples also has revealed significantly reduced RalGAPα2 expression levels in combination with strong increase of inhibitory phosphorylation^35^. Knockout of RalGAPβ increases migration and invasion of PDAC (pancreatic ductal adenocarcinoma) cell lines as well as xenograft growth *in vivo* and accelerates local tumor growth and metastasis^36^. Furthermore, κB-Ras 1/2 deficiency in a KRas^G12D^-driven PDAC mouse model caused faster development of invasive carcinoma and shorter life span of animals^26^. In patients, lower RalGAPα2 expression levels correlate with an invasive phenotype in colitis-associated cancer^31^. They also inversely correlate with survival in prostate^37^ and bladder cancer^34^, and κB-Ras expression levels are reduced in human PDAC tumor samples^25^. These data indicate that RalGAP complexes are important tumor suppressors.

Interestingly, the bi-allelic inactivation of *RALGAPA1* was reported to cause a rare genetic disease leading to impaired neurodevelopment, infantile spasms, muscular hypotonia, and feeding abnormalities^38^, suggesting additional medical relevance of RalGAP beyond cancer.

Here we provide a structural and biochemical characterization of the RalGAP complex and thus a solid framework for understanding the mechanisms underlying its function. RalGAP assembles into a dimer of heterodimers, which results in the formation of an extended 58 nm structure. We demonstrate that this complex architecture is essential for functionality and explain why RGα2 requires RGβ binding to maintain an active conformation. Based on this detailed structural framework, we analyze RalGAP patient variants, suggesting a role in pathological processes such as tumorigenesis.

## Results

### Cryo-EM structure of the RalGAP complex

We established expression of the RGα2/RGβ/κB-Ras2 complex in Expi293F cells and purified it via FLAG affinity purification and size exclusion chromatography, to obtain a monodisperse sample for cryoEM single particle studies (Suppl Fig 1A,B). The sample shows a homogenous set of characteristic extraordinarily long and narrow mustache-shaped particles (Suppl Fig 1C). Already at the initial steps of image processing, it became evident that the 58 nm elongated particles consist of two identical subcomplexes, which are, however, flexibly linked to each other. Therefore, we focused our analysis on the half-particle, which represents the unique portion of the ‘mustache’ structure (Suppl Fig 2, Suppl Table 1). In addition, this focus was necessary for efficient processing, as we required a box size of ∼42 nm for the half-particle alone. To achieve better local resolution, we performed multi body refinement for the core as well as the terminal portions of the ‘half-mustache’. This yielded a map with overall 3.8 Å resolution and 4.4 Å for the focused maps of the hinge region, 3.8 Å for the central region, and 3.9 Å for the peripheral region of the complex (Fig 1B, Suppl Fig 2,3). We were able to unambiguously build an atomic model of the RGα2/RGβ heterodimer using AlphaFold2 models as a starting point (Fig 1C). Several loop regions in both subunits, the very N-terminal portion of RGα2 (residues 1-73) and κB-Ras2, which binds to this region^25^, were not resolved and could not be modeled. The reconstruction includes a small portion of a second RGβ subunit of the adjacent “half-moustache”, which nevertheless allowed us to unambiguously model its N-terminal portion and characterize this interface (Fig 1B,C).

Both RGα2 and RGβ have an overall similar architecture, including an N-terminal α-solenoid HEAT repeat-like domain, with frequent insertions of loops and additional helices within and between HEAT motifs. This region is followed by a globular domain (referred to as stabilization and dimerization (SD) domain in the following) and a GAP domain that is inserted between two helices α^B^ and α^C^ of the SD domain (Fig 1D). The GAP domain of RGα2 folds with a β^1^-α^A^-α^B^-β^2^-β^3^-β^4^-α^C^-β^5^-β^6^-β^7^-β^8^-α^D^ topology that resembles the Asn-thumb GAP domains of TSC2 and RapGAP, with a characteristic catalytic asparagine at position 1742 at the end of helix α^C^ (Suppl Fig 4). RGβ has a similarly folded domain with the core β-sheet structure, but large loops are inserted between β^2^-β^3^ and β^5^-β^6^. Importantly, the catalytic α^C^ helix is substituted by an extended loop (aa 1264-1341) that is partially unresolved in the EM density (aa 1272-1329), rendering the domain a pseudo-GAP (pGAP) lacking the active site residues (Suppl Fig 4).

The SD domains are the central organizing module of the RalGAP complexes and mediate heterodimerization of RGα2 and RGβ in a tail-to-tail fashion, resulting in an overall arch-like shape of the RalGAP heterodimer. In both RalGAP subunits, they show a β^1^-β^2^-β^3^-β^4^-α^A^-β^5^-α^B^-α^C^-α^D^ topology, forming a central 5-stranded β-sheet with two helices on each side (Fig 1D). The β-strands from both SD domains form one continuous sheet to mediate heterodimerization (Fig 1D). In addition, the SD domains interact in trans with the C-terminal HEAT repeats of their binding partner. Interestingly, between β4 and αA of both RGα2 and RGβ SD domains, large loop regions with little secondary structure except for two helical segments are inserted. These loops reach over from RGα2 to RGβ and from RGβ to RGα2, respectively, thereby linking the subunits with each other and accounting for one third of the heterodimer interface (buried surface RGα2-RGβ 10,443 Å², RGα2:RGβ^hug^ 3923 Å², RGα2^hug^:RGβ 3574 Å²) (Fig 1 E,F). We therefore termed these structural motifs ‘hug’ domains and ‘hug’ helices.

The general domain organization of RGα and RGβ is reminiscent of the TSC2 subunit of the tuberous sclerosis complex (TSC) protein complex (Suppl Fig 5), suggesting that RGα, RGβ and TSC2 are likely derived from a common ancestor. However, while TSC2 forms a homodimer via its SD domain, RGα2 and RGβ form a heterodimer (Fig 1D, Suppl Fig 5). There are also differences in the structure of the RalGAP and TSC2 SD domains. The SD domain of TSC2 forms homodimers via a shorter β-sheet and has multiple additional helices inserted between the β-strands, which together form a domain termed ‘saddle’^22^ that stabilizes binding of the TSC1 subunit. These distinct features of the SD domains are required to realize the different complex assemblies and architectures of the TSC and RalGAP complexes.

### Modelling of RalA binding

For a better understanding of the GAP activity mechanism, we further modeled the interaction of RalA with the full RalGAP complex. We first generated an AlphaFold 3 prediction^39^ of RalGAPα2 in complex with RalA and GTP and positioned Ral on the experimental structure by superposing the RGα2 GAP domains of model and prediction (Suppl Fig 6, Fig 2A). The GAP domain of the predicted Ral/RalGAP2 model superimposes well with the GAP domain of the homologous Rap1A/Rap1GAP structure (PDB: 3BRW^14^) with an RMSD of 1.128 Å over 115 Cα atoms. The catalytic asparagine residue (N1742) is properly positioned 4.4 Å away from the γ-phosphate to coordinate a nucleophilic water molecule for stimulation of GTP hydrolysis (Fig 2B). Rap and Rheb have a conserved tyrosine residue in their switch I region that was shown to support GAP-stimulated GTP hydrolysis by stabilizing the γ-phosphate of GTP^14,15,40^. Tyrosine 43 of RalA occupies a similar position (Fig 2B) and thus likely fulfills the same function as in the homologous GTPases. In addition to the GAP domain, the SD domain of RGα2 also makes contacts with the GTPase similar to how the TSC complex is proposed to bind Rheb^22^ (Fig 2A). This interface likely contributes to affinity of the interaction and specificity of GTPase recognition by their cognate GAPs.

**Figure 2:**
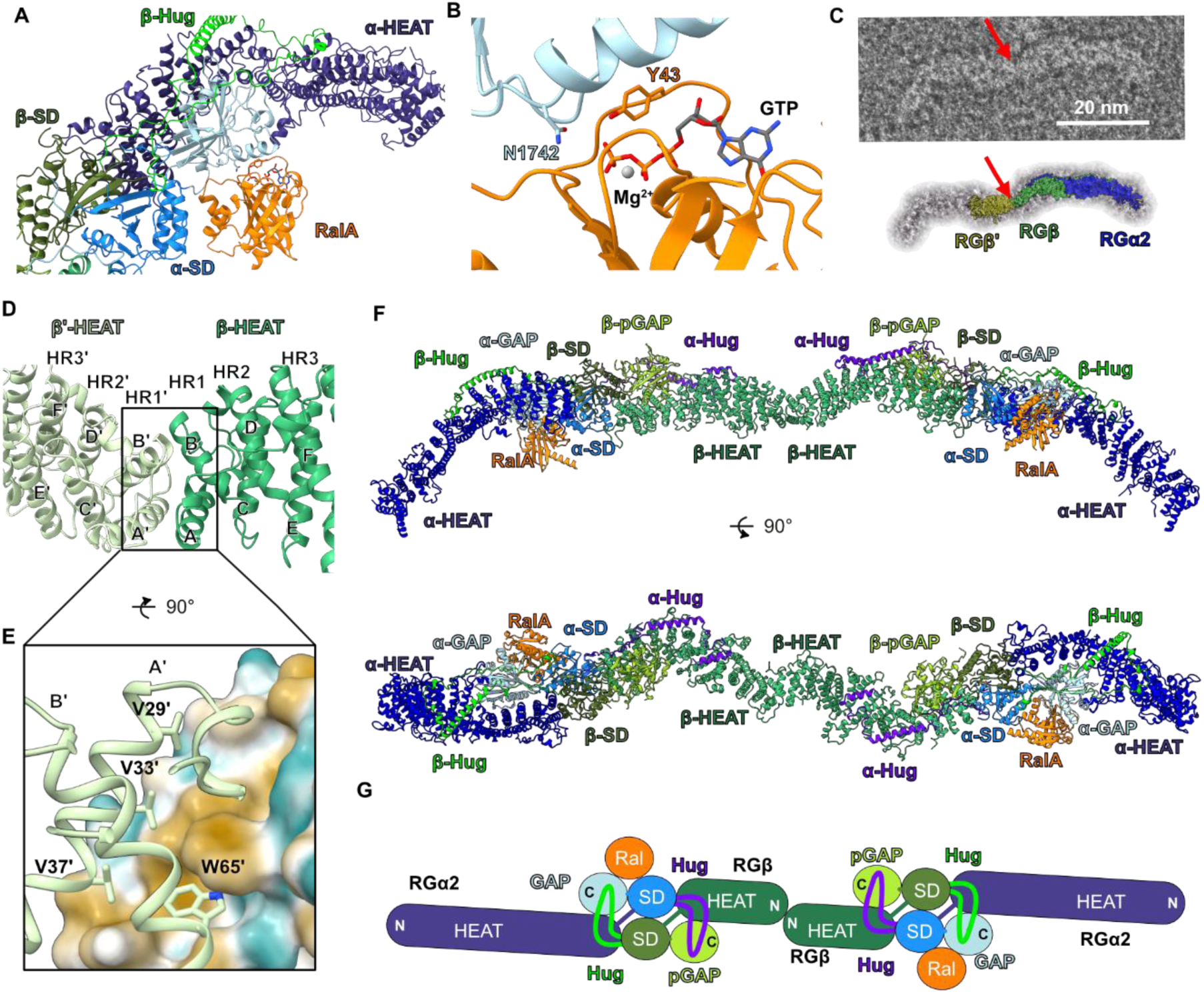
RalA binding and tetramerization of RalGAP. (A) Model of GTP-bound RalA associated with RGα2 based on an AlphaFold 3 prediction. (B) Close-up of the active site. The catalytically relevant amino acids RGα2^N1742^ and Ral^Y43^ are positioned as the equivalent residues in the structure of the Rap-RapGAP complex. (C) Negative stain image of a full RalGAP particle in comparison to the 3D reconstruction of the half-particle. The red arrow marks the connection between two heterodimeric half-particles. (D) N-terminal homodimerization of RGβ (E) Close-up of the homodimerization interface of the RGβ N-terminal domain. One RGβ subunit is shown in surface representation colored according to polarity (orange: hydrophobic, cyan: hydrophilic), the other RGβ is shown as cartoon with key interface residues shown as sticks and labeled. (F) Model of the RalGAP tetramer with RalA obtained by superposition of a second RGα2/RGβ heterodimer on the N-terminal HEAT domain of RGβ’ in the experimental model and the AF3 prediction of RalA interaction with RGα2 GAP domain. (G) Schematic representation of the Ral-RalGAP assembly.

### Tetramerization of RalGAP

Unexpectedly, the structure reveals homodimerization of RGβ via its N-terminal region at the central interface between the two half-particles (Fig 2C). Thus, RGβ mediates the interaction of two RGα2/RGβ heterodimers and leads to the formation of the tetrameric ‘mustache’-shaped RalGAP complex. The RGβ subunits interact via hydrophobic residues in the first two helices of the RGβ HEAT domain (Fig 2D,E). The key amino acids of this interface include a central tryptophane (W65) and a series of valines in the first helix (V29, V33, V37). The small portion of the second RGβ’ subunit visible in the experimental map (Fig 1B) provides sufficient detail to extend the model of RalGAP by superposition of a RGα2/RGβ heterodimer with the RGβ’ N-terminal domain and experimental images of the full particle. This allowed us to create a complete model of the extended, ∼580 Å long RalGAP tetramer. Combined with the modelling of the GAP domain interacting with RalA, we provide a model of the complete RalGAP2 complex engaged with its substrate (Fig 2F,G).

### The unique hug domain of RGβ contributes to catalytic activity through stabilizing effects

Surprisingly, while RGβ is essential for GAP activity in cells and *in vitro*^10^, it does not interact with RGα2 at the active site and - based on the high confidence AlphaFold3 model and in homology to RapGAP - is also unlikely to interact with Ral (Fig 2A). However, the RGβ hug domain makes extensive contacts with RGα2, in particular the long hug helix that runs alongside the HEAT and GAP domains and holds them together (Fig 1F). We suspected that the β-hug domain thus may be indirectly involved in GAP activity. To first test the relevance of the hug domains for complex stability, we generated deletion constructs RGα2^Δhug^ (lacking aa 1498-1593) and RGβ^Δhug^ (lacking aa 1004-1099) lacking these domains. In co-immunoprecipitation experiments, RGα2^Δhug^ still bound full-length RGβ, while RGβ^Δhug^ almost completely lost its ability to associate with RGα2 (Fig 3A), revealing an essential role of RGβ^hug^ in complex assembly. Expectedly, deletion of the hug domain in both subunits abolished complex formation as well. Furthermore, we found that the RGβ hug domain alone was sufficient to form a complex with RGα2 (Fig 3B). These findings suggest that the hug domain of RGβ is crucial for the integrity of the RalGAP complex.

**Figure 3:**
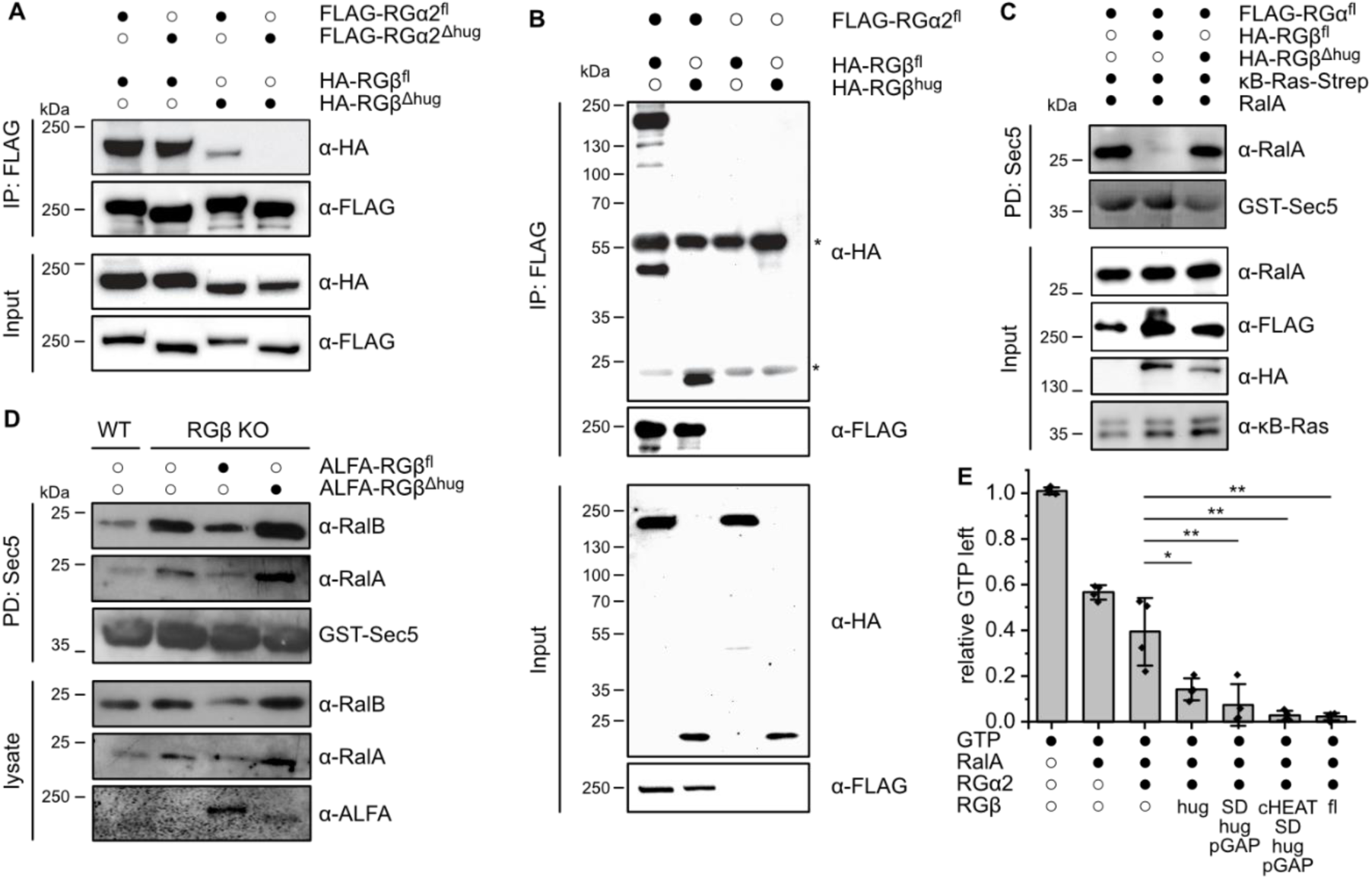
The RGβ hug domain is required for RGα2 GAP activity. (A) Co-immunoprecipitations of RGα2 and RGβ full-length (RGα2^fl^, RGβ^fl^) and hug domain deletion constructs (RGα2^Δhug^, RGβ^Δhug^) from transiently transfected HEK293T cells. (B) Co-immunoprecipitations of RGα2 with RGβ full-length and a construct only comprising the β-hug domain (RGβ^hug^, aa 973-1118) from transiently transfected HEK293T cells. (C) Sec5 pull-down from HEK293T cells transfected with RalA and RGα2 without or with RGβ^fl^ and RGβ^Δhug^. (D) Sec5 pull-down analysis of RGβ knock-out MEFs reconstituted with RGβ^fl^ or RGβ^Δhug^. (E) *In vitro* GAP assay of RalA GTP hydrolysis by RGα2 without or with various RGβ truncation constructs. The same molarity of RGα2 was used in each assay and the activity was normalized to the amount of RGβ present in the individual reactions, n= 4 Two-sample t-test *: p < 0.05, ** p<0.01.

For a functional validation, we analyzed the importance of the RGβ hug domain with a cellular assay where RalA and RGα2 were co-expressed with the different RGβ constructs. The levels of active RalA in these cells were determined by a pull-down assay with the Sec5 Ral effector domain that selectively binds GTP-bound Ral GTPases. RGα2 and RGβ together were able to sustain RalGAP activity, but not the co-expression of RGα2 with RGβ^Δhug^ (Fig 3C). We then used RGβ knock-out mouse embryonic fibroblasts (MEFs), which show dramatically increased levels of active Ral, for reconstitution experiments. Reintroduction of full-length RGβ, but not RGβ^Δhug^, was able to reduce Ral activity in these cells (Fig 3D). To identify a minimal active RalGAP complex, we employed an HPLC-based *in vitro* GAP assay that measures GTP hydrolysis by RalA in the presence of RalGAP (Fig 3E). As previously reported, robust stimulation of GTP hydrolysis was observed for the full-length RGα2/RGβ complex, but not for RGα2 alone. We then co-expressed and purified RGα2 with fragments of RGβ comprising only the hug domain (RGβ^hug^, aa 973-1118), the C-terminal SD domain with hug and pseudo-GAP domain (RGβ^SD-hug-pGAP^, aa 881-1495) and a slightly longer construct that additionally contains the C-terminal portion of the HEAT repeat domain (RGβ^cHEAT-SD-hug-pGAP^, aa 576-1494). Although these constructs were less stable than full-length RGβ, they could be co-purified with RGα2 and these subcomplexes caused robust acceleration of GTP hydrolysis by RalA (Fig 3E). This identifies the hug domain of RGβ as the key element sufficient to render RGα2 catalytically active. Taken together, the structural and biochemical data suggest that the function of RGβ in the enzymatic mechanism of the RalGAP complexes is to stabilize the GAP domain via its hug domain.

### Formation of RalGAP tetramers via RGβ dimerization is required for complex function

We next asked if homodimerization via the N-terminal domain of RGβ is functionally relevant. In co-immunoprecipitations of differently tagged versions of RGβ we observed that indeed, two RGβ copies can interact in cells (Fig 4A). We then introduced mutations of key residues identified in the observed homodimer interface of RGβ (Fig 2D,E). A single point mutant RGβ^D1^ (W65R), a triple dimer interface mutant RGβ^D3^ (V29E, V33Q, V37E), and a combination of both RGβ^D4^ all lost the ability to homodimerize (Fig 4A), while interaction with RGα2 was not affected (Suppl Fig 7A). Co-purification of RGβ^D4^ with RGα2 yielded smaller complexes compared to wildtype as assessed by size exclusion chromatography (Suppl Fig 7B) and negative stain EM analysis (median 580 Å and 289 Å), respectively, (Fig 4B, Suppl Fig 7C,D). RGα2/RGβ^D4^ complexes showed stimulation of GTP hydrolysis *in vitro* comparable to wild-type RalGAP, demonstrating that catalytic activity is not affected by the mutations (Fig 4C). Surprisingly, RGβ^D4^ expression in RGβ KO cells was not able to rescue the phenotype but resulted in unchanged high levels of GTP-bound Ral (Fig 4D). These data demonstrate that an intact tetrameric RalGAP assembly is required for its physiological function, possibly by ensuring proper regulation or localization.

**Figure 4:**
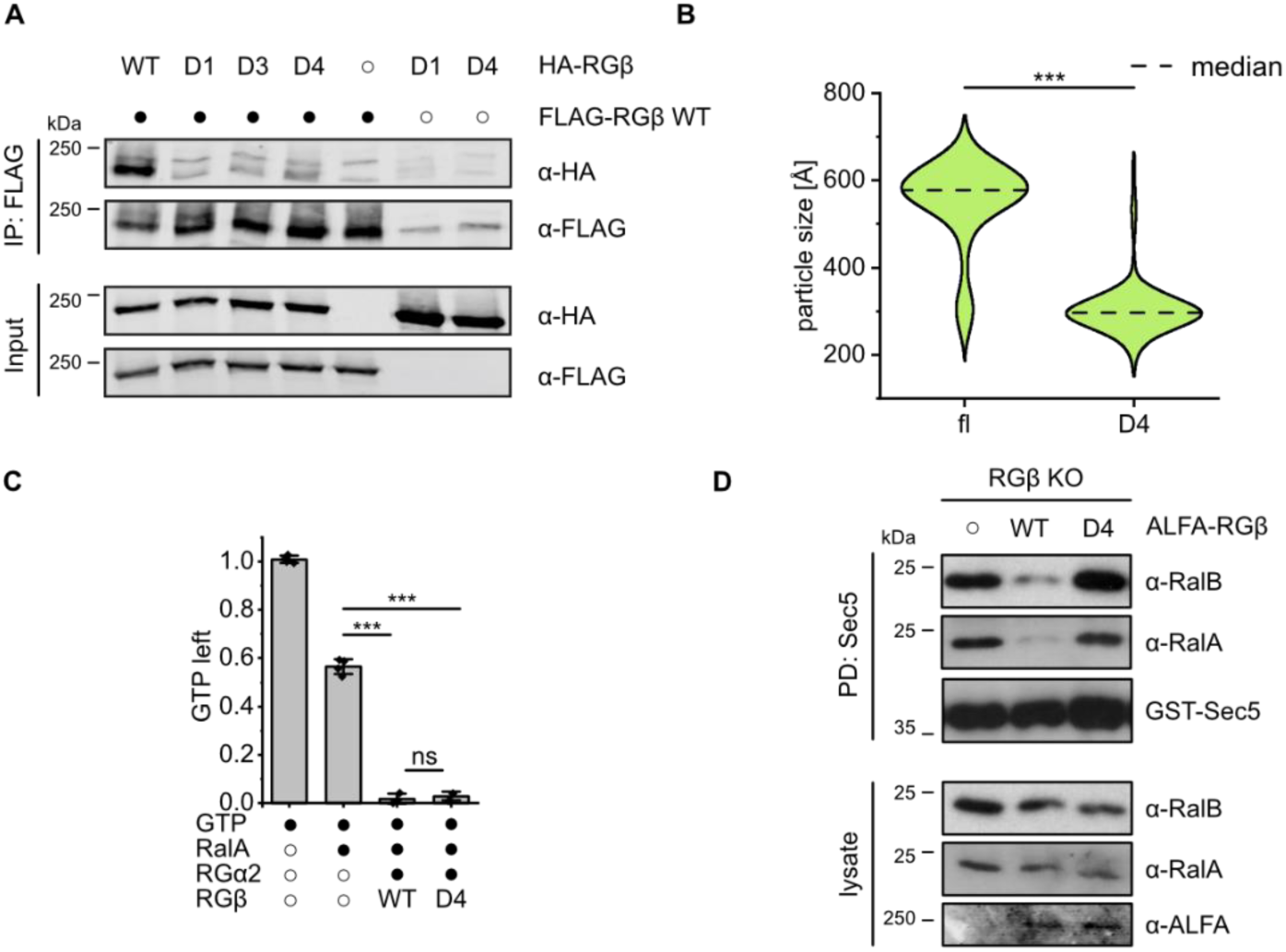
Tetramerization is essential for RalGAP function. (A) Co-immunoprecipitations of differently tagged RGβ full-length (RGβ^fl^) and RGβ variants from transiently transfected HEK293FT cells. WT: wild-type; D1: W65R; D3: V29E, V33Q, V37E; D4: V29E, V33Q, V37E, W65R. (B) Particle length distribution measured by negative stain EM analysis of purified RGα2/RGβ^WT^ and RGα2/RGβ^D4^ complexes. Statistical analysis: Mann-Whitney-test, ***: p<0.001. (C) *In vitro* GAP assay of RalA GTP hydrolysis by RGα2 with RGβ^WT^ or RGβ^D4^. n= 3-4-Statistical analysis: Two sample t-test, ns: p > 0.05, ***: p< 0.001. (D) Sec5 pull-down analysis of RGβ knock-out MEFs reconstituted with RGβ^WT^ or RGβ^D4^.

### Structural analysis suggests pathogenicity of RalGAP variants identified in cancer patients

To provide structural insights into the role of RalGAP complexes as tumor suppressors, we searched for missense mutations in the *RALGAPA1*, *RALGAPA2* and *RALGAPB* genes in cancer patients. In the TCGA PanCancer Atlas Studies (32 studies, 10967 samples), RalGAP coding genes showed the highest missense mutation frequency in uterine and skin cancers (16% of all samples) (Fig 5A). We compiled a list of missense mutations in *RALGAP* genes from uterine and skin cancer patients reported on cBioportal^41,42^, which are classified as variants of uncertain clinical significance (VUS). Their locations are spread over the entire length of the three genes and no hot spot mutations can be identified, which is reminiscent of mutations in *TSC* genes that cause tuberous sclerosis complex (TSC)^43–45^ (Suppl Fig 8A-C). We chose all positions that are reported mutated more than once in the *RALGAPA2* and *RALGAPB* genes for closer inspection. Comparison to the AlphaMissense database^46^ yielded only a small portion of variants to be categorized as likely pathogenic (20%, Suppl Table 2). However, AlphaMissense analyses the isolated proteins and thus primarily classifies variants as likely pathogenic that are expected to directly affect protein folding and/or stability based on their predicted structure. Manual inspection of these variants on basis of the experimental model confirms the assessment that the mutations are likely to negatively affect protein structure (Fig 5B). AlphaMissense analyses proteins in isolation, but a large portion of variants classified as likely benign by AlphaMissense maps to protein-protein interfaces that are revealed by our complex structure (Fig 5B). This includes the Ral binding site, the heterodimerization interface and the hug domains or hug binding interfaces. Distortions of these interfaces are likely to interfere with complex function, again indicating potential pathogenicity of the reported variants.

**Figure 5:**
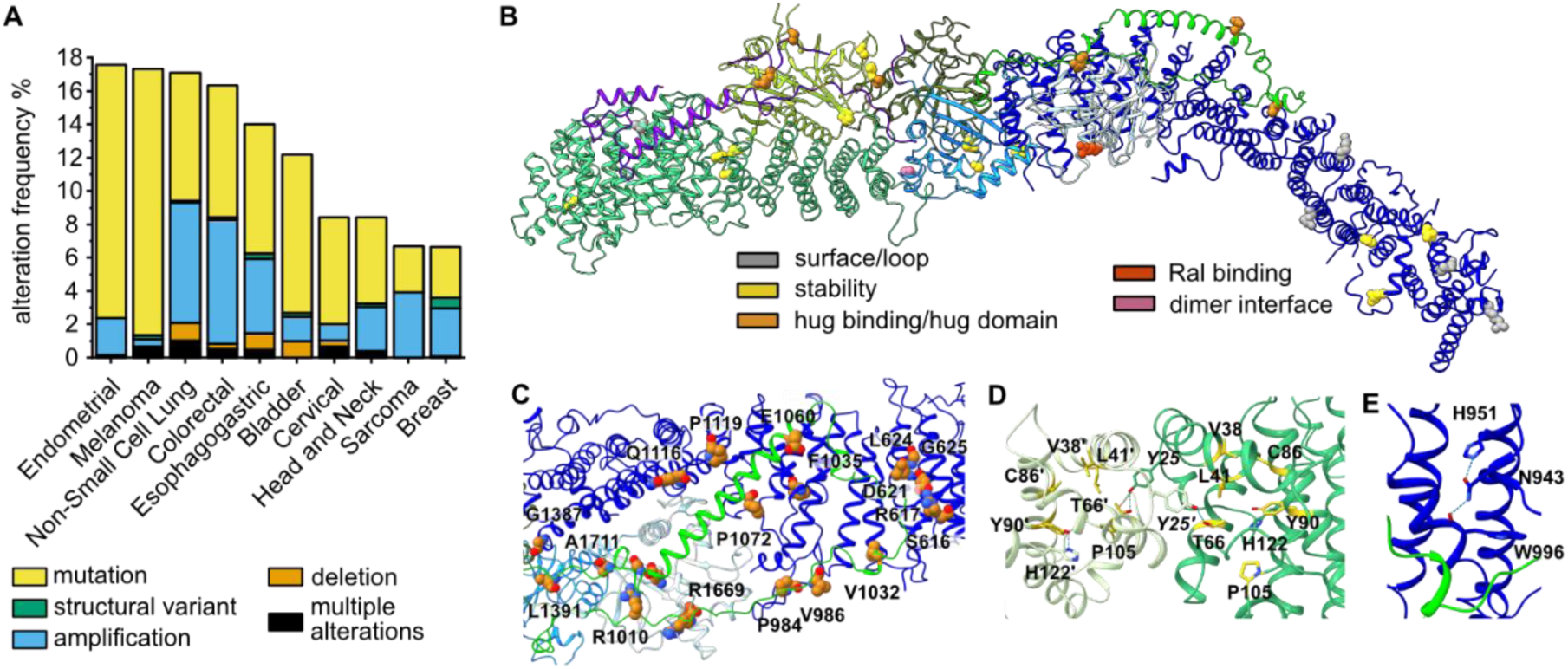
Structure-based assessment of RALGAP variants. (A) Summary of alterations in the *RALGAPA1*, *RALGAPA2* and *RALGAPB* genes reported in the TCGA PanCancer Atlas Studies. The ten cancer types with the highest alteration frequency are shown. (B) Mapping and structure-based classification of *RALGAPA2* and *RALGAPB* variants reported more than once in uterine and skin cancer patients. (C) Close-up of the β-hug/RGα2 binding site with the positions of patient variants shown in sphere representation. (D) Representation of the RGβ N-terminal homodimerization interface with the amino acids at positions of patient variants shown as sticks. (E) Stabilization of the RGβ^hug^ binding site of RGα2 by N943.

We extended our analysis to all variants reported in cancer patients that map to the two essential interfaces - β-hug binding to RGα2 and RGβ N-terminal homodimerization - that we validated by biochemical and cell biological studies. Patient variants are frequently found in both the β-hug domain and the corresponding binding interface on RGα2 (Fig 5C). Furthermore, a cluster of patient variants is found at the RGβ homodimerization interface (Fig 5D). Thus, the analysis of patient variants in the light of the RalGAP structure and its functional characterization suggests that at least some of the reported variants may be pathogenic and could contribute to tumorigenesis. Interestingly, a few variants cluster in a large disordered loop of the RGα2 HEAT domain (672-911), close to the reported AKT phosphorylation sites S696 and T715^27^ (Suppl Fig 8, Suppl Table 2). These variants may therefore be pathogenic without affecting RalGAP structure.

Independent of somatic mutations, bi-allelic loss of RALGAPA1 was described as cause for the genetic disease NEDHRIT (neurodevelopmental disorder with hypotonia, neonatal respiratory insufficiency, and thermodysregulation)^38^. Interestingly, one patient carried an allele with a pathogenic missense mutation (c.3227A>G, p.Asn1076Ser). This position is conserved between RGα1 and RGα2, and the corresponding residues (N1075 of RGα1 and N943 of RGα2) are involved in stabilizing the HEAT repeat domain at the β-hug binding site of the protein (Fig 5E). Mutation of RGα2 N943 (and likely RGα1 N1076) is thus expected to lead to a destabilization of RGα and impaired complex formation, demonstrating that already a mild structural destabilization of RalGAP can be disease-causing.

## Discussion

The structural analysis of RalGAP reveals a unique tetrameric complex architecture. RGα and RGβ form a tail-to-tail heterodimer via their SD domains, and two heterodimers interact via head-to-head dimerization at the N-terminal domain of RGβ. The complex contains two active sites at the GAP domains of RGα. Thus, RalGAP remarkably differs in its structure from the TSC complex, although the subunits RGα, RGβ and TSC2 show a conserved domain architecture and likely are derived from a common ancestral gene. In the TSC complex, TSC2 forms homodimers via the SD domains and binds the regulatory TSC1 subunit. TSC1 shows no similarity to RGα or RGβ and interacts with a long dimeric coiled-coil domain alongside the TSC2 homodimer and thus introduces asymmetry. TSC1 is not required to promote activity of the TSC2 GAP domain per se as a minimal TSC2 construct was reported that has comparable activity on Rheb as the full-length TSC complex *in vitro*. Instead, TSC1 is mainly responsible for the proper localization and function of the TSC complex in cells, by binding phosphoinositides, WIPI proteins and TBC1D7^16,47–49^.

Our study elucidates the relevance of RGβ for RalGAP function, identifying two relevant molecular mechanisms. First, the hug domain of RGβ binds RGα2 and thereby enables the GAP domain of RGα2 to adopt its active conformation. Presumably, RGα2 does not fold properly in the absence of β-hug, suggesting that the RalGAP subunits need to undergo a mandatory swap of their hug domains to complete their structure. Second, RGβ is required for the formation of the RalGAP tetramer, which does not influence catalytic activity *in vitro* but functionality in cells. At this stage, we can only speculate why tetramerization is important, but it is likely a prerequisite for proper spatiotemporal control. This highlights how accurately RalGAP needs to be regulated in a physiological setting.

A previous study that mapped interaction sites within the RalGAP complexes identified a short peptide sequence in RGβ (‘β-blockatide’, residues E1125-T1162 in the RGβ sequence used in this study) that was able to bind RGα1 and reduce RalGAP complex formation and thereby protein stability^50^. We find that in the structure of RalGAP this peptide is part of the SD domain of RGβ but does not directly contact RGα2 (Suppl Fig 9). The effects observed with β-blockatide are thus likely caused by interference with RGβ folding rather than direct inhibition of subunit interactions.

Along these lines, it is interesting to note that many of the variants of uncertain clinical significance (VUS) in the *RALGAP* genes reported in cancer patients are likely to negatively affect complex function based on our structural assessment. The sheer number of variants makes a systematic experimental assessment hardly feasible. On the other hand, the automated computational categorization of variants by AlphaMissenese is currently not sufficiently reliable as it does not account for protein structure and function in the context of the assembled protein complex. The individual analysis of variants based on the experimental structure of RalGAP, we report here, thus provides a valuable tool for an initial assessment of VUSs.

Interestingly, no obvious mutation clusters can be identified but variants are spread over the entire lengths of all three *RALGAP* genes. This is reminiscent of TSC, where pathological variants are also found throughout both *TSC1* and *TSC2* without apparent hot spot mutations. It is also interesting to note that just like in TSC, where mutations occur in the GAP complex but rarely in the cognate GTPase Rheb, no oncogenic Ral mutations are reported. Potentially, Ralopathies may also rather be caused by loss-of-function of the corresponding GAP protein.

While loss or reduction of RalGAP subunit expression has been shown to promote tumorigenesis in human cancer cell lines and mouse models and has been described to occur in a subset of pancreatic cancer patients^1^, a function of RalGAP in uterine or skin cancer or the impact of RalGAP mutations in general has so far not been addressed. We report that a significant number of patient variants are expected to destabilize the complex or map to the functional important region that we newly identified, like the β-hug domain or the RGβ homodimerization site. It thus seems worthwhile to more closely investigate (mutated) RalGAP as a tumor suppressor relevant for different human cancers. Our study provides the necessary structural framework towards a better understanding of the relevance of RalGAP in cancer biology.

## Methods

### RalGAP Protein Purification

DNA sequences encoding for RalGAPα2 and RalGAPβ were cloned into a p3xFLAG-CVM7.1 and pKH3 vector, respectively. κB-Ras2 DNA sequence was cloned into pCTAPa vector^51^. RalGAP complexes were expressed in Expi293F cells. For a 60 ml culture, total amount of 60 µg plasmid DNA encoding the respective RalGAP subunits were mixed with 1500 µL PBS and incubated with PEI (1:3 DNA:PEI) for 15 min at room temperature before adding to 150 x 10^6^ cells in a 250 mL flask. After incubating cells for 4 h at 37 °C and 8 % CO_2_, 3.5 µM valproic acid was added and the culture filled up to a final volume of 60 mL with Expi293 medium (Thermo Fisher Scientific). The cells were harvested after 3-4 days, and the pellets snap frozen in liquid nitrogen.

Cell pellets were thawed on ice and resuspended in 12 mL lysis buffer F (for full length constructs: 50 mM Tris-HCl pH 7.5 150 mM NaCl, 1 mM MgCl_2_, 5 % Glycerol, 1% Triton-X100) or lysis buffer C (for minimal GAP constructs: 50 mM Tris-HCl pH 7.5 150 mM NaCl, 4 mM MgCl_2_, 5 % Glycerol, 0.3 % CHAPS) supplemented with 1:50 protease inhibitor mix HP (Serva) and 0.5 µM DTT. Cells were incubated on a rotating wheel at 5 rpm, 4 °C for 20 min and debris removed by centrifugation (12,000 x g, 45 min; 4 °C) and filtration (0.45 µM; cellulose acetate syringe filter). Clear lysate was passed four times over M2 α-flag beads (Sigma-Aldirch) equilibrated in lysis buffer. Beads were washed twice with 2 ml lysis buffer and twice with 2 ml RGH buffer (20 mM HEPES pH 7.5, 150 mM NaCl, 2 mM MgCl_2_) and subsequently eluted with 0.1 µg/µl 3xFLAG peptide (ApexBio A6001 or TargetMol TP1274) in RGH and 1 mM TCEP were added. The combined elution fractions were concentrated with an Amicon Ultra Centrifugal filter (Millipore). For use in GAP assays, buffer was exchanged to RGA buffer (30 mM HEPES pH 7.5, 150 mM NaCl). For preparation of cryo-EM samples, the concentrated protein was loaded on a Superose 6 increase 5/150 column (Cytiva) equilibrated with RalGAP-EM buffer (20 mM HEPES pH 7.5, 150 mM NaCl, 2 mM MgCl_2_, 1 mM TCEP). The peak fractions were directly used for grid preparation.

Murine RalA G-domain (aa 9-183) was amplified via PCR and inserted into pCDF6p vector via BamHI/NotI restriction sites. *Escherichia coli* BL21 cells were transformed with this construct and grown in TB media to an OD_600nm_ of 0.9-1.0. Expression of the GST-tagged protein was induced by adding IPTG (0.5 mM) and cells were grown overnight at 16 °C. Cell pellets from a 2 l culture were resuspended in buffer L (50 mM NaH_2_PO_4_ pH 8.0, 300 mM NaCl, 1 mM MgCl_2_, 1 mM DTT, 5 % glycerol, supplemented with lysozyme, DNaseI, protease inhibitor mix HP (Serva)) and lysed (Microfluidizer). The lysate was cleared by centrifugation (39 000 x*g*, 45 min, 4 °C) and loaded onto equilibrated GSH-Agarose beads (Serva) three times and washed three times with 20 ml buffer L. The protein was cleaved from the beads with PreScission protease overnight and loaded onto a Superdex 75 pg 16/600 HiLoad size exclusion column (GE) equilibrated with buffer R (30 mM HEPES pH 7.5, 150 mM NaCl, 10 % glycerol). Peak fractions were pooled, concentrated to 800 µM and stored at -70 °C.

### Negative stain grid preparation

Grids were glow discharged and incubated with 4 µl of protein solution for 2 min. Excess of protein solution was blotted away (Whartman paper No 1) and the grid washed twice with 10 µl water, once with 10 µL uranyl acetate (UA) and incubated with UA for 45 sec before staining solution was blotted away. Grids were analyzed using a Talos L120C G2 TEM operating at an acceleration voltage of 120 kV at 120,000x magnification.

### Sample vitrification and cryo-EM data acquisition

For preparing the grids for cryo-EM, 4 μl sample at a concentration of 0.7 mg/ml were applied onto a freshly glow-discharged QuantiFoil 2/1 Cu300. The solution was incubated for 2 min, then manually blotted to remove excess liquid, and another 4 µl of protein solution was added, together with 1 µl 0.018% Triton-X100 solution. The excess liquid was then immediately blotted followed by vitrification in liquid ethane using a Vitrobot II automatic plunge-freezer (Thermo Fisher Scientific).

### Data acquisition

Datasets were acquired with a 300 kV Titan Krios G4 microscope (Thermo Fisher Scientific) equipped with an E-CFEG, a Selectris X energy filter and a Falcon 4i direct electron detector operated by the software EPU (Thermo Fisher Scientific). A total of 64,606 micrographs in five sub-datasets at 0°, 30° and 42° stage tilt angles were collected in Electron Event Representation mode (EER) at a nominal magnification of 215k, corresponding to a pixel size of 0.58 Å/px. Most micrographs were collected at defocus of -0.5 to -4.2 μm. The Selectris X energy filter was used for zero-loss filtration with an energy width of 10 eV. A total dose of ∼60 e-/Å2 was aimed for by adjusting the exposure time of each sub-dataset to an appropriate value (between 2.7 and 3.6 seconds per micrograph). The details of dataset collection are summarized in Suppl Table 1.

### Image processing and 3D reconstruction

EER movies were motion corrected in RELION5^52^ using its own Motioncorr2^53^-like algorithm. CTF estimation was performed using CTFFIND4^54^, and particles were selected using crYOLO^55^. The initial particle pick using the generalized model of crYOLO was suboptimal due to the elongated and partially overlapping particles. To improve the picking model and thereby achieve a reasonable selection of particles, Relion5 was used to perform multiple rounds of 2D classification, initial model generation, 3D-refinement and re-extraction with improved centering, followed by retraining and repicking in crYOLO using the improved picking models.

With the thus optimized picking model, 5,519,589 particles were extracted from the motion-corrected micrographs by crYOLO, with a window size of 720 × 720 pixels, and binned to a box size of 360x360 pixels (1.16 Å/pixel). This binning factor was kept throughout further refinement. A subset of 887,720 particles were selected by 2D classification, keeping only classes that showed high-resolution features and complete particles which most likely were picked at somewhat similar positions along the particle. The latter was important as the elongated and relatively featureless mustache-like complex allowed origin offsets of up to several hundred pixels, thereby complicating subsequent 3D-refinements. 3D-classification into four classes resulted in only two approximately equally populated classes and two classes with only 37 and 252 particles, respectively. The better resolved 3D class containing 420,975 particles was selected for further processing. 3D-refinement reached 4.1 Å resolution and could be improved to 3.98 Å after CTF and aberration refinement and another round of 3D-refinement in Relion5. Bayesian polishing and another round of CTF and aberration refinement, each followed by 3D auto-refine, further improved the resolution to 3.83 Å and a strongly improved map interpretability, which declined towards the box edges, as can be expected due to the elongated particle with dimensions exceeding the chosen box dimensions. We therefore performed a final round of multi-body refinement as implemented in Relion5, splitting the particles into three bodies, one in the center of the box, and two following the particle towards the box edges. This resulted in three partial maps with further improved map interpretability, and resolutions ranging from 3.79 to 4.35 Å.

### Model building

A model of the RGα2/RGβ complex was calculated using AlphaFold2^56^ running on a local high performance computing cluster (PALMA II), docked into map from 3D reconstruction with ChimeraX^57^ and energy optimized with ISOLDE^58^. In an iterative process, the model was optimized using real space refinement in Phenix Refine^58^ against the full density map, and manual model building with Coot^59^ using the full density map as well as the resampled multi body refinement maps. To generate a composite map of the three partial maps, these maps were resampled onto the initial full map in ChimeraX and combined by using the combine_focused_map tool from the Phenix suite.

AlphaFold3^39^ was used to generate a model of RGα2 in complex with RalA, GTP, and Mg^2+^. The model was superimposed with the experimental structure by aligning the GAP domains and the position of the Ral GTPase was combined accordingly with the experimental model. The energy of a composite model of the RalGAP core with Ral was minimized using the YASARA forcefield^60^.

### Visualization and analysis of cryoEM maps and models

Structure visualization and analysis was done with ChimeraX (UCSF)^57,61^. Local resolution gradients within a map were calculated with RELION5 and visualized with ChimeraX. 3D angular distribution plots were generated in Relion5. 2D histograms of the angular distribution were generated using angdist^62^. The 3D Fourier shell correlation of cryo-EM maps was calculated using the remote 3DFSC processing server^63^.

### GTPase activity assay

0.3 µM GAP proteins were incubated with 10 µM mouse RalA G-domain (aa 9-183) in assay buffer (30 mM HEPES, 150 mM NaCl, pH 7.5) supplemented with 20 mM EDTA and 1 mM DTT. 5 mM MgCl_2_ and 50 µM GTP were added to start the reaction. The sample was briefly mixed and aliquots were snap frozen in liquid nitrogen at given time points and samples stored at – 70 °C. Nucleotide ratios were analyzed by HPLC. The assay samples were boiled for 5 min at 95 °C and precipitated protein was removed by centrifugation. Of the cleared supernatant, 10 µl were injected in an Agilent 1260 Infinity II HPLC system equipped with an autosampler and a DAD HS detector. Analytes were separated on an AMAZE HA mixed phase column (Helix Chromatography). The separation of G-nucleotides was achieved by a stepwise double gradient with increasing the buffer (200 mM KH_2_PO_4_ pH 2.0, 30-80 %) and acetonitrile (15-20 %) concentrations. The UV traces at 254 nm were used to monitor nucleotide elution. UV traces were semi-automatically analyzed with OriginPro 2024 (OriginLab Corporation). Traces were base line corrected, the peaks corresponding to GDP and GTP integrated and the portion of GTP calculated. The remaining GTP amount was normalized to GTP at t = 0 h. In case of sub-stoichiometric complexes, the RGβ/RGα2 ratio was calculated from Coomassie stained SDS-PAGE band intensities and the remaining GTP value compensated by this factor. Individual biological repeats were calculated from three technical repeats.

### Cell lines

HEK293FT (RRID:CVCL_6911) cells were obtained from Thermo Fisher Scientific (R70007). Mouse embryonic fibroblasts (MEFs) were generated from a C57BL/6 mouse embryo at day E12.5 and SV40-immortalized^25^. CRISPR/Cas9 mediated knockout of RalGAPβ was performed as described below^64^. All cell lines were cultured in DMEM high glucose (Sigma-Aldrich, D6546) supplemented with 10 % FBS (Biowest, S1810), 10 mM L-Glutamine (Sigma-Aldrich, G7513) and 50 U Penicillin-Streptomycin (Thermo Fisher Scientific, 15070063) at 37 °C with 5 % CO_2_.

### CRISPR/Cas9-based generation of RalGAPβ knockout MEFs

For CRISPR knockouts sgRNA sequences were chosen with the CRISPick tool^65,66^ (https://portals.broadinstitut.org/gppx/crispick/public) and primers designed with BbsI compatible overhangs. Annealed Oligos were cloned into px458 or px459 plasmids (gifts from Feng Zhang; Addgene plasmids # 48138 and # 62988^64^). SV40-immortalized WT MEFs were simultaneously transfected with px458 and px459 plasmids containing sgRNAs targeting upstream (RGβsg1 agaagcagtagtggtagtgt) and downstream (RGβsg2: gctgctaactccagttgcag) of exon 2 of RalGAPβ using jetPRIME® DNA and siRNA transfection reagent (Polyplus, 101000046). 24 h after transfection GFP-positive cells were FACS-sorted and returned to culture to recover overnight. Cells were then selected for 36 hours with 2 µg/ml puromycin. Single cell clones were obtained by limited dilution. Successful deletion of RGβ was tested using PCR screening (mRGβ screen 2F tgaaagggaaatgtcggaaa, mRGβ screen 2R tgagttcctgccttggtttt) and confirmed by RT-qPCR using primers targeting inside the deleted exon (mRGβRTEx2_F cagtggctggtagtgagagt, mRGβRTEx2_R gcaacaccaaagccataatcc) and immunoblot.

### Reconstitution of RalGAPβ knockout MEFs

N-terminally ALFA-tagged RGβ^fl^, RGβ^D4^ or RGβ^Δhug^ were cloned into a modified pITR-TTP vector ^47^ via MluI and NotI restriction sites. RGβKO MEFs were transfected with pITR, pITR-ALFA-RGβ^fl^, pITR-ALFA-RGβ^D4^ or pITR-ALFA-RGβ^Δhug^ and the transposase expressing pCMV-Trp plasmid (9:1 ratio) using jetPRIME DNA and siRNA transfection reagent (Polyplus, 101000046) for stable reconstitution. Cells were selected for 48 hours with 2 µg/ml puromycin and maintained with 1.5 µg/ml puromycin afterwards. To confirm exogenous expression, proteins were isolated with Bäuerle lysis buffer (20 mM Tris pH 8, 350 mM NaCl, 20 % glycerine, 1 mM MgCl_2_, 0.5 mM EDTA, 0.1 mM EGTA, 1 % NP-40 supplemented with 1 mM DTT, Protease and Phosphatase Inhibitor Cocktail (Thermo Fisher Scientific, A32965, A32957) and analyzed by immunoblot.

### Co-Immunoprecipitation

HEK293FT cells were seeded to 10 cm petri dishes and grown to 70 % confluency. The medium was exchanged to starvation medium and prepared DNA incubated for 4 h at 37 °C 5 % CO_2_. Medium was changed to growth medium. The next day, medium was removed and the cells washed 1-2 times with ice cold PBS. 1 ml CoIP buffer (40 mM HEPES, 120 mM NaCl, 10 mM MgCl_2_, 0.3 % CHAPS, pH 7.4) supplemented with PIC (1:100) was added to the cells and cells collected using a gum wiper. Cells were incubated for 20 min on a stirring wheel at 4 °C before debris was removed by centrifugation. The clear lysate was taken and incubated with 20 µl 3xflag beads slurry (1:1 in lysis buffer) for 2-4 h. Beads were pelleted, the supernatant removed, and the beads washed three times with CoIP buffer. Beads were resuspended in 1 x SDS-LD and analyzed by western blotting.

### Ral effector pull down

HEK293FT cells were transiently transfected as described previously. Cells were lysed in 800 ml SLB (50 mM Tris-HCl, 100 mM NaCl, 4 mM MgCl2, 1 % Triton-X100, pH 7.5 at 4 °C). The Ral-binding domain of rSec5 was purified from *E. coli* as GST-fusion construct and loaded onto GSH beads. 20 µg GST-rSec5 were added to the cleared cell lysate and incubated (45 min, 4 °C, 5 rpm). The supernatant was removed and the beads washed with SLB three times before resuspending in 1x SDS loading dye.

### Immunoblot

Proteins separated by SDS-PAGE (sodium dodecyl sulfate polyacrylamide gel electrophoresis) were blotted on Immobilon-FL (Millipore, IPFL00010) membrane using a semidry system (Trans-Blot Turbo Transfer System, 1704150, Bio-Rad). For imaging on the Odyssey CLx Imaging System (LI-COR), membranes were blocked for 1 h in Blocking Buffer (0.1 % casein (Sigma-Aldrich, E0789) in 0.2 x PBS). Primary antibodies were diluted 1:1000 in 1:1 PBS:Blocking Buffer with 0.1 % Tween and incubated with the membranes at 4 °C overnight. Secondary antibodies were applied 1:4000 for 1 h in 1:1 PBS:Blocking Buffer with 0.1 % Tween and 0.01 % SDS: IRDye 800CW donkey anti-mouse (LI-COR, 926-32212) or IRDye 680RD donkey anti-rabbit (LI-COR, 926-68073). For chemiluminescent imaging, membranes were blocked for 1 h in 3 % BSA (Serva Electrophoresis, 11930) in TBS-T (1 % Tween). Primary antibodies were diluted 1:1000 in 1.5 % BSA/TBS-T and incubated with the membranes at 4 °C overnight. Secondary antibodies were applied 1:4000 for 1 h in 1 % dry milk powder (Carl Roth, T145.2) in TBS-T. Secondary antibodies used were Horseradish Peroxidase goat anti-mouse (Jackson ImmunoResearch, 115-035-044) or Horseradish Peroxidase goat anti-rabbit (Jackson ImmunoResearch, 111-035-045). For imaging on the ChemoSTAR Touch (intas) system, membranes were in blocked for 1 h in 5 % non-fat dried milk (Applichem) in 1 x TBS. Primary antibodies were diluted 1:7000 (anti-flag), 1:2000 (anti-HA) and 1:100 (anti-RalA) in TBS and incubated with the membranes at 4 °C overnight. Membranes were washed three times with TBS/T before HRP-coupled secondary antibodies were applied 1:25000 in TBS and incubated for 2 h at RT (Polyclonal rabbit anti-mouse, Dako P0260; polyclonal swine anti-rabbit, Dako P0217). Membranes were washed and chemiluminescence induced with the SuperSignal West Pico PLUS detection reagent kit (Thermo Scientific, Ref 34577).

The following primary antibodies were used: anti-RalA (Proteintech, 13629-1-AP), anti-RalB (OTI2C4, Origene, TA505880), anti-ALFA (Nanotag, N1582), anti-FLAG (clone M2, Sigma-Aldrich, F1804), anti-HA (clone C29F4, Cell Signaling Technology, 3742), anti-HA (clone 16B12, Biolegend) and anti-κB-Ras (provided by S. Ghosh, Columbia University^25^).

### Statistical Analysis

Data curation and statistical analysis were done with OriginPro 2024 (OriginLab Corporation). Normality was tested with a Shapiro-Wilk test. Significance was tested with two-sample t-test for normally distributed samples and Mann-Whitney test for non-normally distributed samples.

## Author contributions

DK designed the project. AO, CG and DK supervised the study. RR purified proteins and prepared EM samples with MC and BUK. BUK collected and processed cryo-EM data with RR. RR performed structural modeling. RR, LA, and EM performed functional assays. RR, AO, CG and DK wrote the manuscript. All authors discussed the results, commented on the draft and approved the final version of the manuscript.

## Acknowledgements

This work was supported by grants from Deutsche Forschungsgemeinschaft (DFG, German Research Foundation) to DK (KU2531/2, KU2531/6) and AO (OE531/4-1). The cryo-EM data were collected at “Cryo-EM SoN”, the cryo-EM infrastructure of the University of Münster, funded by the DFG (project number 496113311). The cryo-EM data processing and AlphaFold2 modeling was carried out on the Palma II HPC (DFG INST 211/667-1) of the University of Münster. We thank Mark Nellist for helpful discussions.

## Notes

### Competing Interest Statement

The authors have declared no competing interest.

