## Supplementary Information for "Structure and mechanism of the RalGAP tumor suppressor complex"

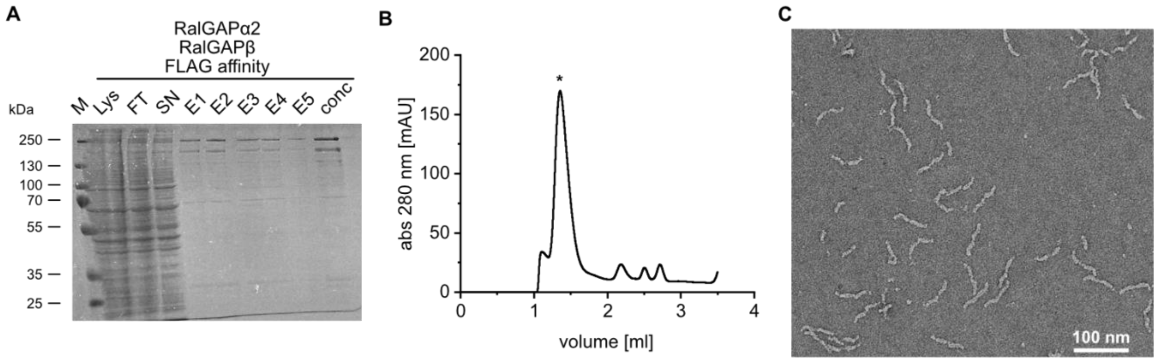

**Supplementary figure 1: Purification of RalGAP. A** Coomassie stained SDS-PAGE gel of a RalGAP FLAG affinity purification. **B** Size exclusion chromatography profile of RalGAP (Superose 6 increase 5/150 column (Cytiva), 20 mM HEPES pH 7.5, 150 mM NaCl, 2 mM $MgCl_2$, 1 mM TCEP). Peak fraction used for EM analysis is marked by an asterisk. **C** Representative negative stain EM micrograph of RalGAP preparation.

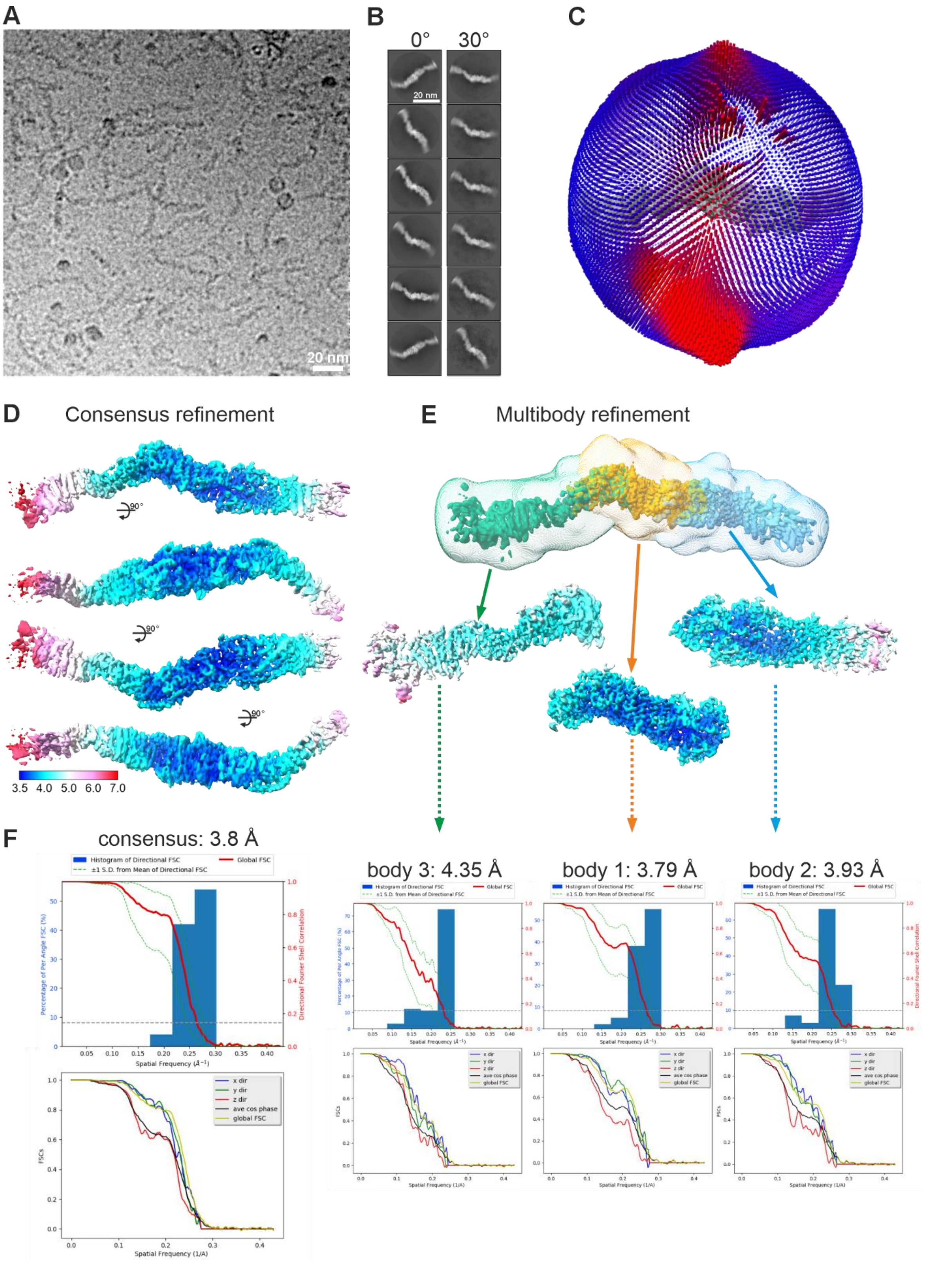

**Supplementary figure 2: Cryo-EM analysis of RalGAP. A** Representative cryo-EM micrograph. **B** Representative reference-free 2D class averages from datasets collected at 0° and 30°, respectively. **C** Angular distribution of RalGAP particles in the final consensus refinement. **D** Density maps of the RalGAP consensus refinement colored by local resolution prior multi-body refinement. **E** The used masks for Relion5 multibody-refinement and the generated locally refined maps colored by local resolution. **F** 3D-FSCs of the consensus and masked maps.

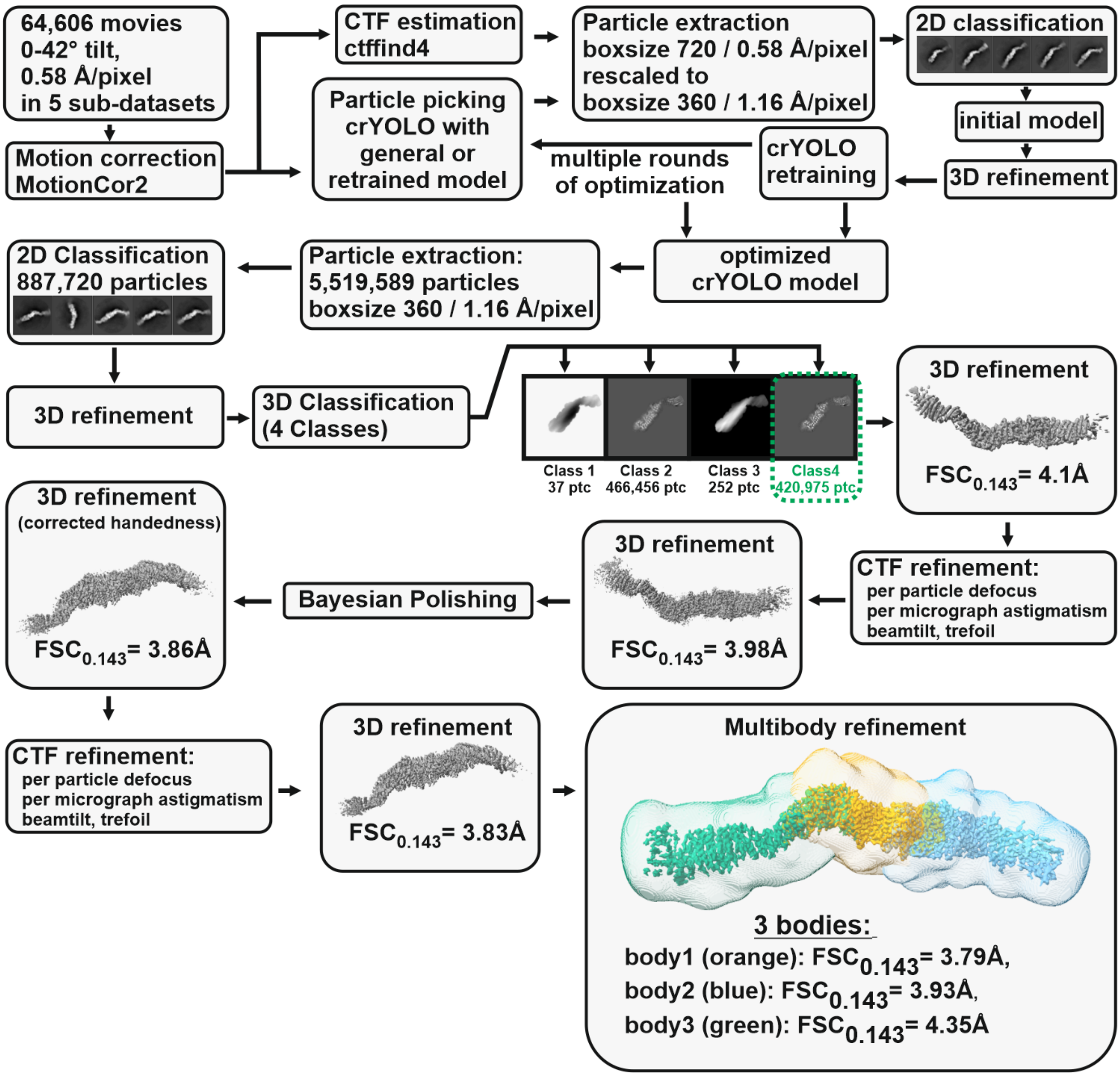

**Supplementary figure 3: Cryo-EM processing workflow for structure determination of RalGAP.**

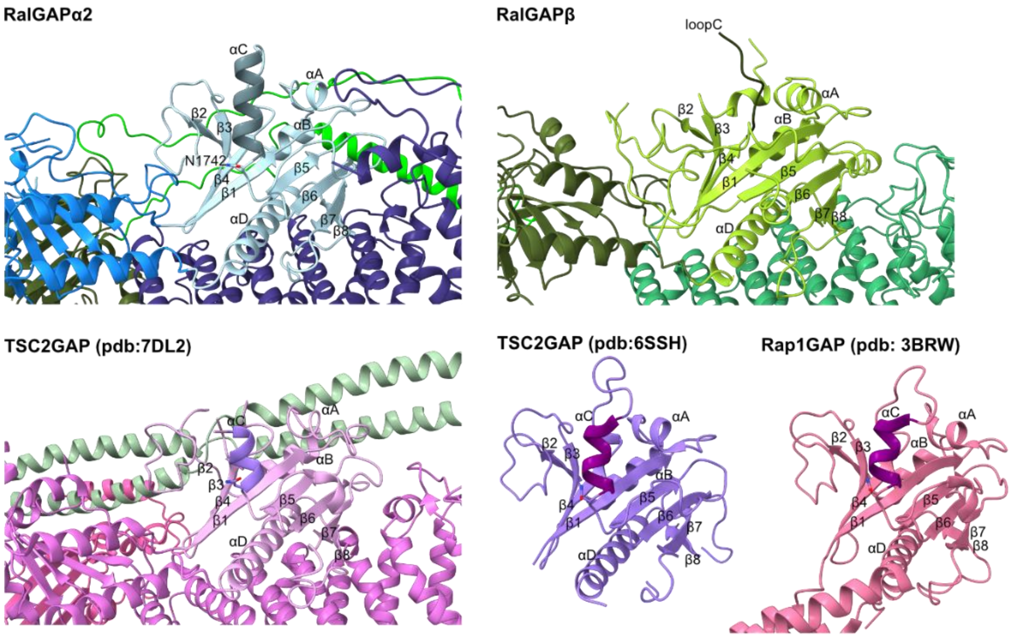

**Supplementary figure 4: Comparison of Asn-thumb GAP domains.** Experimental structures of Asn-Thumb GAP domains shown in the same orientation: RGα2 and RGβ of the RalGAP (this study), human TSC2 in the TSC complex[1], isolated TSC2 GAP domain of *Chaetomium thermophilum*[2], Rap1GAP[3]. The catalytic helices αC are highlighted in darker color, except for RGβ that does not contain a helix αC.

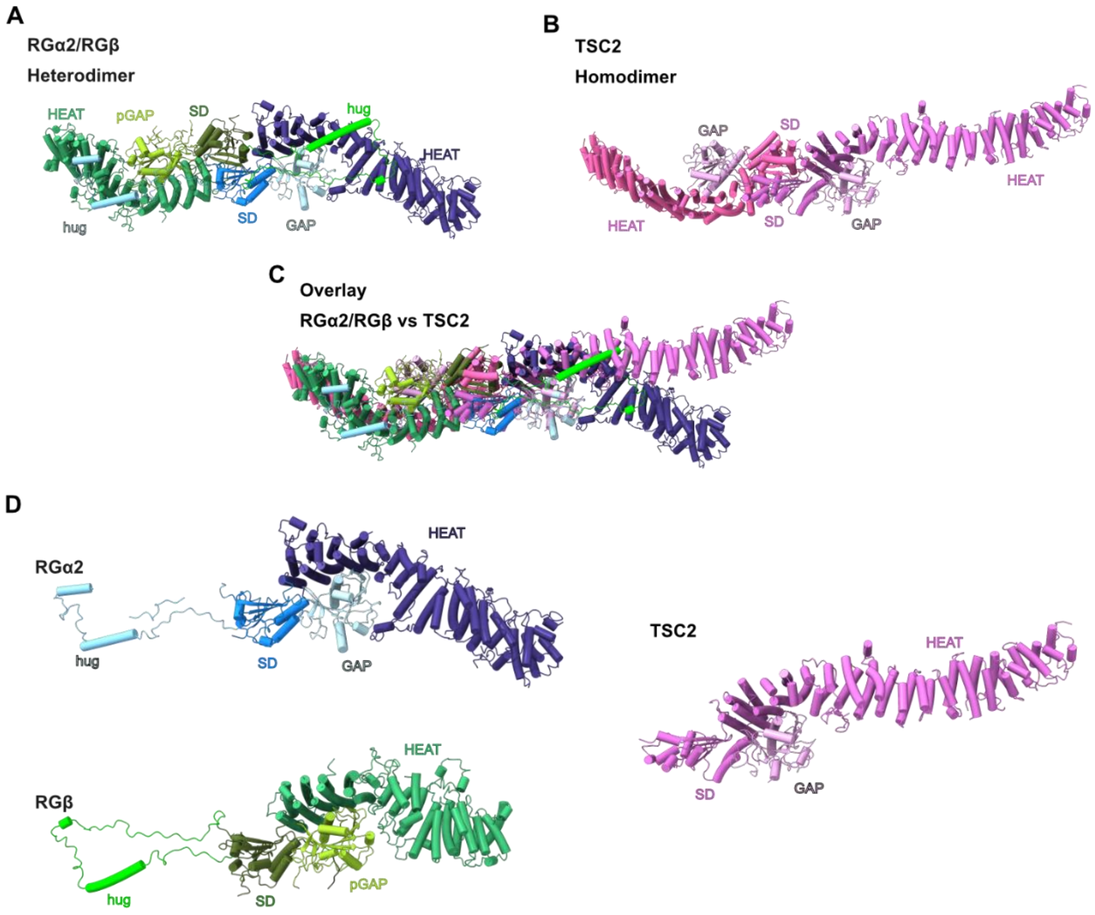

**Supplementary figure 5: Structural comparison of RalGAP and the TSC complex. A** Structure of the heterodimeric RalGAP half-particle. **B** Structure of the TSC complex. **C** Superposition of the RalGAP and TSC complexes. **D** Structures of RGα2, RGβ and TSC2 with the SD and (p)GAP domains in the same orientation.

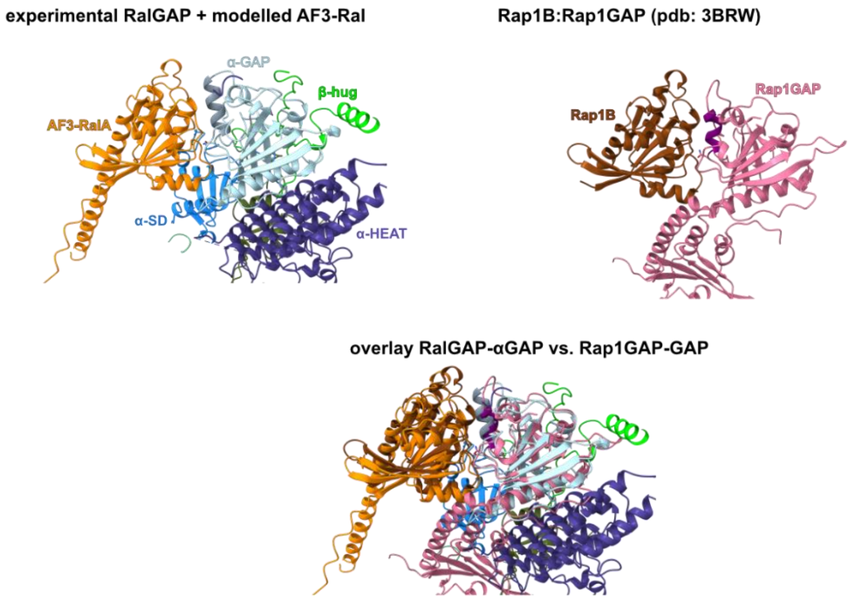

**Supplementary figure 6: Modeling of the interaction between Ral and RalGAP.**

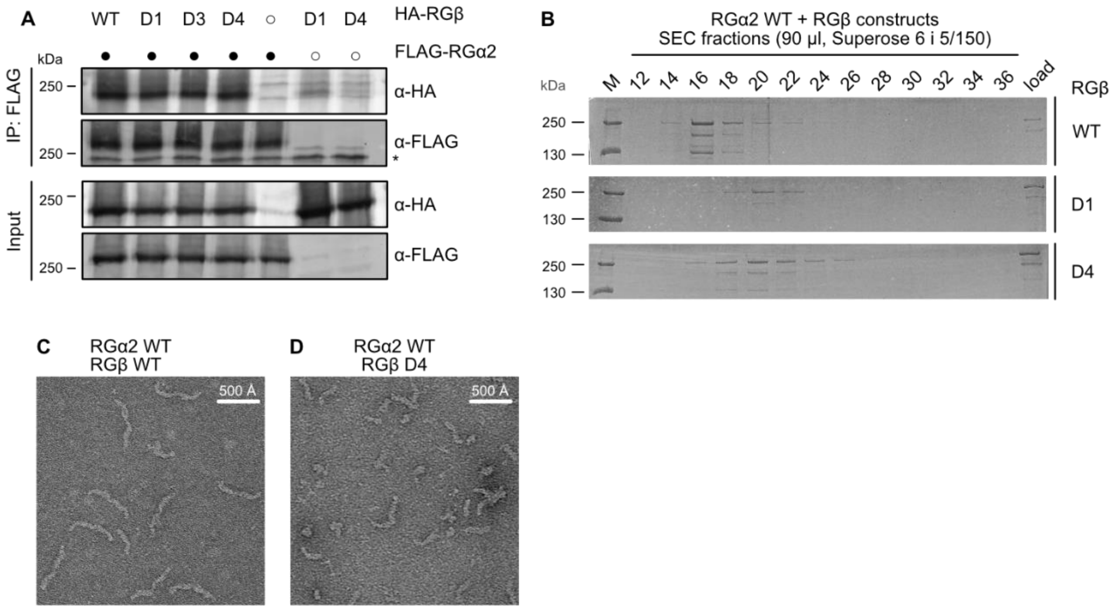

**Supplementary figure 7: Homodimerization of RGβ. A** Co-immunoprecipitations of RGα2 and RGβ variants from transiently transfected HEK293FT cells. D1: W65R; D3: V29E, V33Q, V37E; D4: V29E, V33Q, V37, W65R; WT: wild-type. **B** Analysis of RalGAP complexes with RGβ WT and RGβ mutants with size exclusion chromatography (Superose 6 increase 5/150 column (Cytiva), 20 mM HEPES pH 7.5, 150 mM NaCl, 2 mM $MgCl_2$, 1 mM TCEP). **C, D** Representative negative stain images of purified RGα2/RGβ WT and D4 mutant.

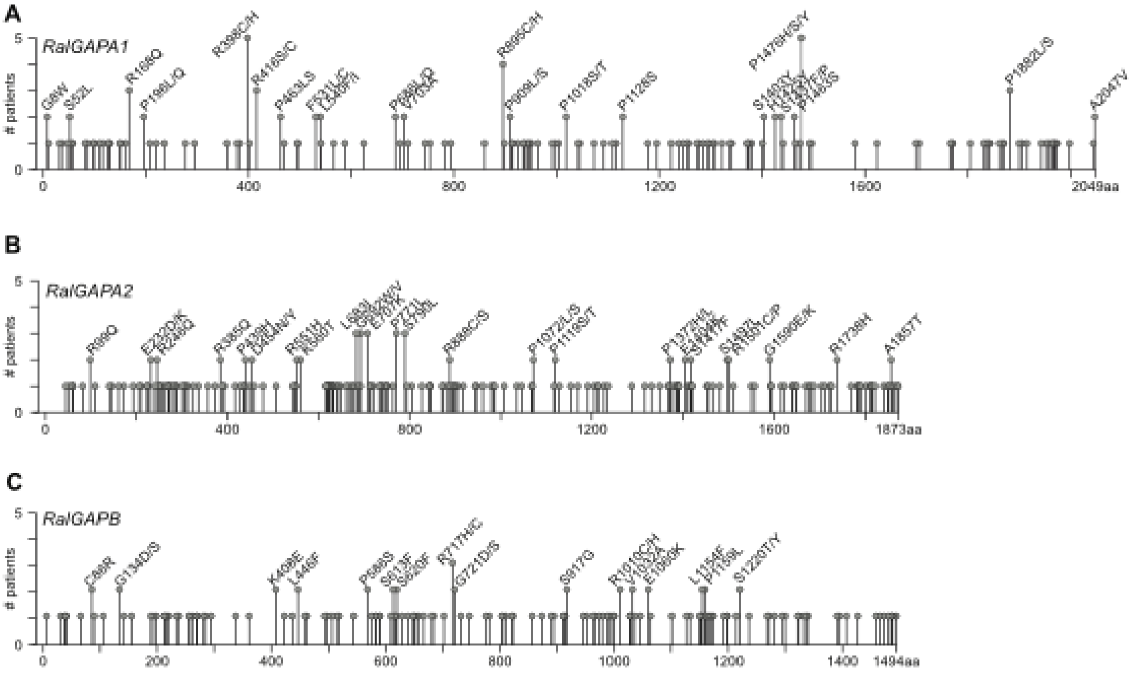

**Supplementary figure 8: Distribution of *RALGAP* variants reported in cancer patients.** Lollipop plots of variants of the **A** *RALGAPA1*, **B** *RALGAPA2*, and **C** *RALGAPB* genes reported in samples form patients with uterine and skin cancer. Generated with cBioportal[4,5] and modified.

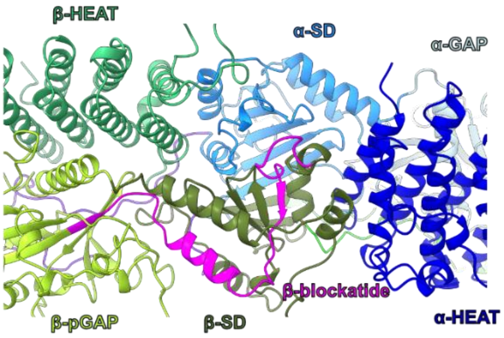

**Supplementary figure 9: Mapping of the β-blockatide peptide in the RalGAP structure.**

**Supplementary table 1: Cryo-EM data collection and refinement statistics of RalGAP.**

| | | | |
|---|---|---|---|
| **Data collection** | | | |
| Microscope | Titan Krios G4 (Selectris X, E-CFEG) | | |
| Voltage (kV) | 300 | | |
| Camera | Falcon 4i | | |
| Pixel size (Å) | 0.58 | | |
| Tilt angle | all Micrographs | picked Micrographs | Particles initial/final |
| 0° | 7,239 | 5,935 | 719,364 / 90,076 |
| 0° | 4,136 | 3,978 | 381,698 / 63,172 |
| 30° | 3,770 | 3,231 | 304,681 / 40,685 |
| 0° | 40,001 | 27,246 | 3,814,505 / 214,791 |
| 42° | 9,445 | 2,733 | 299,341 / 12,251 |
| total number of particles | 5,519,589 | | |
| Number of frames | 793-819 | | |
| Number of fractions | 61-63 | | |
| Total electron dose ($e^-$/Å$^2$) | 60 | | |
| Defocus range (µm) | -0.5 – -4.2 | | |
| **Atomic model composition** | | | |
| Chains | 3 | | |
| Symmetry imposed | C1 | | |
| Non-hydrogen (protein) atoms | 24089 | | |
| Residues | 3028 | | |
| particle substack | 420,975 | | |
| Ligand atoms | - | | |
| **Refinement (Phenix)** | Composite map | | |
| RMSD bond (Å) (# > 4σ) | 0.004 (0) | | |
| RMSD angle (°) (# > 4σ) | 0.978 (4) | | |
| Model to map fit, CC mask | 0.81 | | |
| Model to map fit, CC box | 0.78 | | |
| Resolution (FSC@0.143, Å) | 3.3 (masked)<br>3.6 (unmasked | | |
| B-factor (mean, Å$^2$) | 104.10 | | |
| **Validation** | | | |
| Clashscore | 7.67 | | |
| Ramachandran outliers (%) | 0.00 | | |
| Ramachandran allowed (%) | 2.81 | | |
| Ramachandran favoured (%) | 97.19 | | |
| Molprobity score | 1.84 | | |
| EMRinger score | 1.322 | | |

**Supplementary table 2: RGα2 and RGβ variants from cBioportal.** Missense variants reported more than once in uterine and skin cancer patients are listed with the AlphaMissense classification and a structural assessment based on the experimental cryo-EM structure of the RalGAP complex.

| Variant | AlphaMissense pathogenicity score | | Structural assessment |
|---|---|---|---|
| ***RalGAPα2*** | | | |
| R99Q | 0.2315 | likely benign | Surface |
| E232D | 0.1309 | likely benign | Surface |
| E232K | 0.2572 | likely benign | Surface |
| R246Q | 0.1286 | likely benign | Stability |
| R385Q | 0.0808 | likely benign | Stability |
| P439H | 0.9001 | likely pathogenic | Stability |
| D454N | 0.0736 | likely benign | Loop |
| D454Y | 0.0972 | likely benign | Loop |
| R551H | 0.0802 | likely benign | Surface |
| K560T | 0.1399 | likely benign | Surface |
| L683I | 0.0694 | likely benign | Loop |
| G692V | 0.3277 | likely benign | Loop |
| G692W | 0.6096 | likely pathogenic | Loop |
| E707K | 0.4844 | ambigous | Loop |
| P771L | 0.0662 | likely benign | Loop |
| S790L | 0.0502 | likely benign | Loop |
| R888C | 0.0794 | likely benign | Loop |
| R888S | 0.1767 | likely benign | Loop |
| P1072L | 0.3627 | ambigous | β Hug Binding |
| P1072S | 0.3614 | ambigous | β Hug Binding |
| P1119S | 0.1139 | likely benign | β Hug Binding |
| P1372H | 0.658 | likely pathogenic | Stability |
| P1372L | 0.6238 | likely pathogenic | Stability |
| E1404K | 0.8282 | likely pathogenic | Stability |
| S1417F | 0.4812 | ambigous | Stability |
| S1497L | 0.0954 | likely benign | Hug domain |
| R1501C | 0.0667 | likely benign | Hug domain |
| R1501P | 0.0837 | likely benign | Hug domain |
| G1590E | 0.068 | likely benign | Hug domain |
| G1590K | 0.103 | likely benign | Hug domain |
| R1738H | 0.8389 | likely pathogenic | Ral Binding |
| A1857T | 0.2198 | likely benign | Heterodimerization |
| ***RalGAPβ*** | | | |
| C86R | 0.998 | likely pathogenic | Stability |
| G134D | 0.192 | likely benign | Surface |
| G134S | 0.075 | likely benign | Surface |
| K408E | 0.346 | ambigous | Loop |
| L446F | 0.7678 | likely pathogenic | Loop |
| P568S | 0.0751 | likely benign | Stability |
| S613F | 0.6525 | likely pathogenic | Stability |
| S620F | 0.738 | likely pathogenic | Stability |
| R717C | 0.3867 | ambigous | Loop |
| R717H | 0.2507 | likely benign | Loop |
| G721D | 0.7047 | likely pathogenic | Loop |

| | | | |
|---|---|---|---|
| G721S | 0.0924 | likely benign | Loop |
| S917G | 0.236 | likely benign | α Hug binding |
| R1010C | 0.22 | likely benign | Hug Domain |
| R1010H | 0.1713 | likely benign | Hug Domain |
| V1032A | 0.9069 | likely pathogenic | Hug Domain |
| V1032M | 0.9322 | likely pathogenic | Hug Domain |
| E1060K | 0.1533 | likely benign | Hug Domain |
| L1154F | 0.1729 | likely benign | Stability |
| P1159L | 0.358 | ambigous | Stability |
| S1220T | 0.0836 | likely benign | Stability |
| S1220Y | 0.2139 | likely benign | Stability |

**Supplementary references.**
